# Perturbing postural stability during treadmill walking with dysfunctional electrical stimulation

**DOI:** 10.1101/2025.05.20.655058

**Authors:** Tomomi Miyata, Naoto Izumi, Kakeru Kimura, Takeshi Yamaguchi, Shinichiroh Yamamoto, Kei Masani

## Abstract

Functional electrical stimulation is commonly used to enhance human movement through low-level electrical activation of muscles. More recently, dysfunctional electrical stimulation (DFES) has been proposed as a method to perturb gait by artificially inducing discomfort and mimicking inadequate muscle activity. Here we investigated strategies to induce internal perturbations during treadmill walking using DFES by systematically varying the timing and target muscle. Eleven healthy participants walked at three different speeds while DFES was applied to the tibialis anterior (TA), soleus (SOL), rectus femoris (RF), and biceps femoris (BF) muscles at 25%, 50%, 75%, and 100% of the gait cycle, each for a duration of 0.2 seconds. The gait cycle was time-locked to heel contact (0%). Results showed a significant reduction in the anterior-posterior margin of stability compared to baseline, particularly when DFES was applied to the SOL at 75%, the RF at 50%, and the BF at 75% of the gait cycle. Under these conditions, increased knee flexion and shorter stride intervals were observed relative to baseline. In conclusion, we identified effective DFES conditions to induce postural instability during walking. By mimicking inadequate muscle activity, DFES provides a promising method to study dynamic balance control and mechanisms underlying falls in neurological populations.

## Introduction

Falls during walking pose a significant, potentially life-threatening risk, particularly for older adults and individuals with neurological conditions [1–4]. Understanding the mechanisms of falls and recovery responses has been a longstanding focus in movement science. To study gait instability, physical perturbation methods are commonly used, such as floor translations, tether pulls, or changes in surface friction, to apply external forces or torques to the body [5–9]. These methods have proven valuable for evaluating reactive balance, especially in populations with known balance impairments. However, such externally applied perturbations do not reflect internal disturbances like motor impairments or muscle weakness [10,11], which are often cited as primary causes of falls in clinical populations. Thus, while traditional mechanical perturbations can simulate instability, they may fail to replicate the internally driven disruptions that more accurately characterize real-world falls in these groups.

A promising alternative is to introduce internal perturbations that mimic inadequate or inappropriate muscle activity during walking, which are disturbances more representative of those experienced by older adults or individuals with neurological conditions. Functional electrical stimulation (FES), which applies electrical impulses to activate specific muscles and produce functional movements [12–14], has traditionally been used to improve mobility. More recently, however, dysfunctional electrical stimulation (DFES) has been introduced to perturb locomotor control by inducing uncomfortable, unnatural muscle activations during walking. Manczurowsky et al. [15] applied DFES to the right hamstrings at the beginning of the swing phase to create a controlled, uncomfortable disruption of normal muscle coordination. Their goal was to test whether such an internal perturbation would interfere with cold (non-emotional, logical) executive function, hypothesizing shared neural substrates with motor adaptation. While their muscle selection and stimulation timing were based on pilot testing, specific details were not reported. Nonetheless, the study was lauded for its methodological innovation. As Budde and Gronwald commented [16], “The innovative use of DFES in this study represents a crucial advance, allowing researchers to investigate internally generated impairments while reducing confounds from the varied pathophysiological profiles of clinical populations.” This perspective aligns closely with the motivation behind the present study.

In this study, we explore a novel application of FES to generate internal perturbations within the walking control system. Unlike mechanical perturbations, DFES-induced muscle activity disrupts motor coordination without the participant’s voluntary input, mimicking the unexpected muscle activations that may underlie real-world falls. Here, we investigated strategies to systematically induce internal perturbations during treadmill walking by varying both the timing and target muscle of DFES. We further examined the influence of walking speed on postural responses. Our goal was to identify effective DFES conditions for provoking postural instability, with the broader aim of advancing understanding of fall mechanisms and informing future fall-prevention interventions.

## Methods

### Participants

Eleven healthy adults (6 females, 5 males; age: 26.3 ± 3.3 years; height: 172 ± 10 cm; weight: 64.0 ± 8.9 kg) participated in the study. All participants provided written informed consent prior to participation. The study protocol was approved by the institutional ethics committee.

### Experimental Procedure

Participants walked on a treadmill (RO-FORM Carbon T7, ICON Health & Fitness Inc., UT, USA) while wearing a safety harness to prevent falls. Surface electrodes were placed over four muscles of the right leg: the tibialis anterior (TA), soleus (SOL), rectus femoris (RF), and biceps femoris (BF). Reflective markers were affixed bilaterally to the second metatarsal, heel, ankle, knee, pelvis, and shoulder for kinematic tracking.

Participants first walked at 1.1 m/s to determine their maximum tolerable amplitude of FES, delivered via an electrical stimulator (MyndSearch, MyndTec Inc., ON, Canada). Stimulation was triggered by a footswitch placed under the right heel. The maximum tolerable amplitude was used throughout the experiment.

Each participant completed a total of 18 treadmill trials: one baseline trial and five FES trials at each of three walking speeds—1.1 m/s (slow), 1.3 m/s (medium), and 1.5 m/s (fast). These speeds were chosen to cover the range of preferred walking speeds for both older and younger adults as reported in prior studies [17–21]. In the FES trials, stimulation was applied to one of the four target muscles at one of four gait cycle timings—25%, 50%, 75%, or 100%—relative to right heel contact (defined as 0%). Each unique muscle–timing combination represented one of 16 distinct FES conditions. For example, stimulation of the TA at 25% of the gait cycle was labeled “TA25%.” The duration of each stimulation burst was 0.2 seconds.

FES timings were pre-programmed based on typical stride intervals and adjusted for participants whose stride durations varied by more than ±10% from the norm. Each condition was applied 16 times within a trial, with at least eight steps between stimulations. Trials were pre-randomized, and participants walked for 30 seconds in the baseline trial and for 240 seconds in each FES trial. During all trials, participants kept their arms crossed to minimize upper body motion. A ten-minute rest period was provided after every five trials.

Kinematic data were recorded at 200 Hz using a motion capture system (Raptor-E, Motion Analysis Corp., California, USA). Footswitch signals and FES pulse timing were recorded at 2000 Hz using a 16-bit analog-to-digital converter (USB-6255, National Instruments, Texas, USA).

### Data Analyses

All data were processed using MATLAB 2023b (MathWorks, Massachusetts, USA). Joint angles of the hip, knee, and ankle were calculated, along with stride interval, right foot clearance, and the margin of stability (MoS) in the anterior–posterior (AP) direction. MoS was computed using the toe marker as the base of support reference. Given that all targeted muscles primarily influence sagittal plane dynamics, AP MoS was selected as the primary outcome measure for assessing postural stability.

Displacements from the baseline condition were computed for each stride. For MoS, the largest negative displacement in the AP direction was extracted per stride. For joint angles, both the largest positive and negative displacements were used. Heel contact was identified from the footswitch signal, while toe-off was determined based on the AP velocity of the toe marker.

To accommodate cases in which heel contact was absent in post-FES strides, the gait cycle was defined from 10% after the first toe-off to 110% of the subsequent toe-off. The stride during which FES was applied was labeled as Post0, followed by Post1 through Post5. The stride immediately preceding Post0 was labeled Pre1. Strides were excluded from analysis if the FES onset deviated by more than ±10% of the gait cycle from the intended timing (see Table S1).

### Statistical Analyses

Bartlett’s test was used to assess homogeneity of variance. For comparisons of stimulation timing within each muscle condition, one-way analysis of variance (ANOVA) was used when variances were homogeneous. When variance assumptions were not met, Kruskal–Wallis tests were applied. Bonferroni corrections were used for post hoc comparisons, and statistical significance was set at p < 0.05.

To confirm that participants had recovered to baseline postural stability between FES applications, Dunnett’s tests were used to compare all parameters at the Pre1 stride to baseline. No significant differences were observed, confirming stability had returned prior to subsequent stimulation events.

## Results

Although three walking speeds were tested, no significant differences were observed among them. Additionally, all measured parameters returned to baseline levels by Post2 stride, and no clear signs of postural instability were evident beyond this point. Therefore, we report results from Post0 and Post1 strides under the fast walking condition in this section. Results for the medium and slow conditions are provided in the Supplementary Material (Figures S1, S2). For the fast-speed condition, one participant’s data was excluded from the TA condition due to incorrect FES timing, resulting in a sample size of n = 10 for TA, and n = 11 for SOL, RF, and BF conditions.

Figure 1 illustrates example time series of hip, knee, and ankle joint angles, along with MoS, for a representative participant under the SOL75% condition, from Post0 to Post5 strides. Clear reactions to FES were observed in the knee and ankle joints, as well as in MoS at Post0 and Post1.

**Figure 1.**
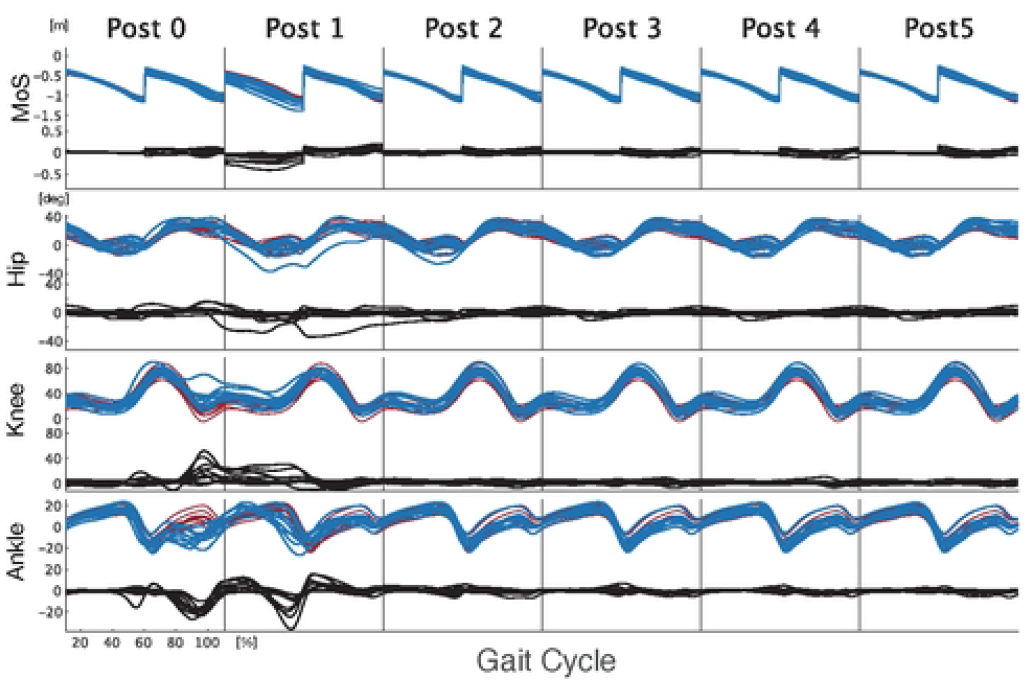
Example time courses of hip, knee, and ankle joint angles and MoS during a FES trial under the SOL75% from Post0 to Post5 strides. For each parameter, the upper blue line represents the baseline trial, the red line represents the FES trial, and the lower black line represents the displacement from the baseline for the respective parameter.

### Joint Angles

Figure 2 shows the largest positive and negative joint angle displacements during Post0 and Post1 strides. At the hip joint, significant deviations from baseline were observed in most conditions (denoted by + or ++ in Figure 2A), with RF50% showing the largest negative displacement at Post0.

**Figure 2.**
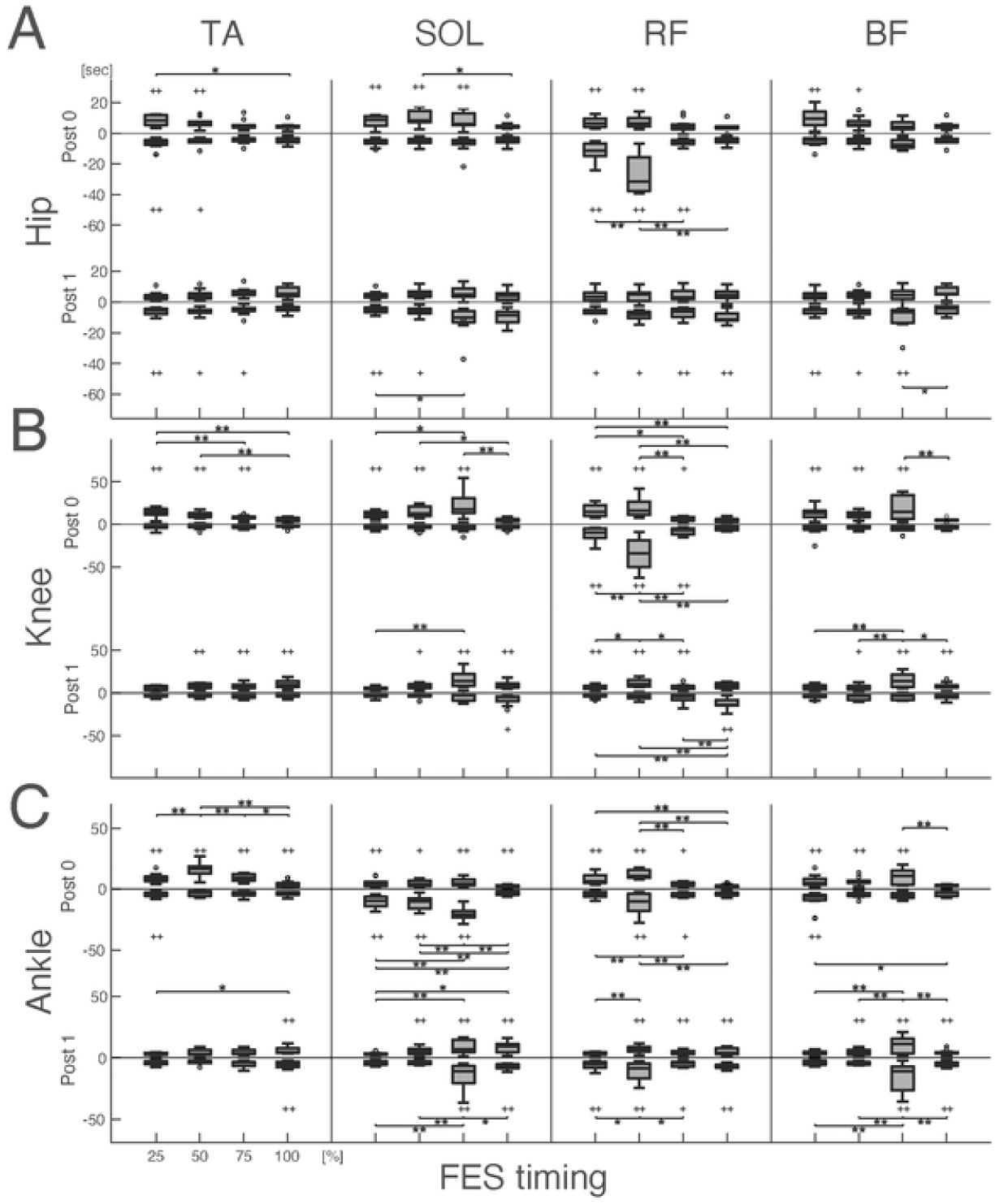
The largest positive and negative displacement value from the baseline in (A) hip joint, (B) knee joint, and (C) ankle joint angle during Post0 and Post1 strides. The + symbol indicates a significant difference from the baseline (+: p < 0.05, ++: p < 0.01), while the * symbol indicates a significant difference between the ones at the timing conditions within each target muscle condition (*: p < 0.05, **: p < 0.01).

For the knee joint, significant positive and/or negative displacements were also observed in nearly all conditions (Figure 2B). At Post0, RF50% induced the greatest positive and negative displacements among the four timing conditions, and was the only condition in which both were significantly greater than baseline within the same stride. At Post1, the BF75% condition showed the largest positive knee displacement. SOL75% also induced high positive displacements at Post0 and Post1, though differences among timing conditions were not significant. These displacements typically occurred just before heel contact, resulting in increased knee flexion at touchdown (as shown in Figure 1). Notably, only RF conditions resulted in significantly greater negative knee displacements at Post0, occurring during FES application.

For the ankle joint, all conditions exhibited significant differences from baseline (Figure 2C). The largest positive displacements at Post0 were observed under TA50% and RF50%, while the largest negative displacements occurred under SOL75% and RF50%. At Post1, positive displacements were greatest under BF75%, and negative displacements under SOL75% and BF75%. These negative values reflected increased plantarflexion at toe-off or reduced dorsiflexion at the end of the swing phase. Importantly, all three joints showed significantly larger positive and/or negative displacements under the SOL75%, RF50%, and BF75% conditions compared to other timing conditions within the same muscle.

### Stride Interval, Foot Clearance, and Margin of Stability (MoS)

Figure 3 summarizes the displacements from baseline in stride interval (Figure 3A), foot clearance (Figure 3B), and MoS (Figure 3C). A significantly prolonged stride interval at Post0 was observed only under RF50%, although no significant differences were found compared to other timing conditions within RF. At Post1, SOL75% and BF75% exhibited significantly shorter stride intervals compared to baseline.

**Figure 3.**
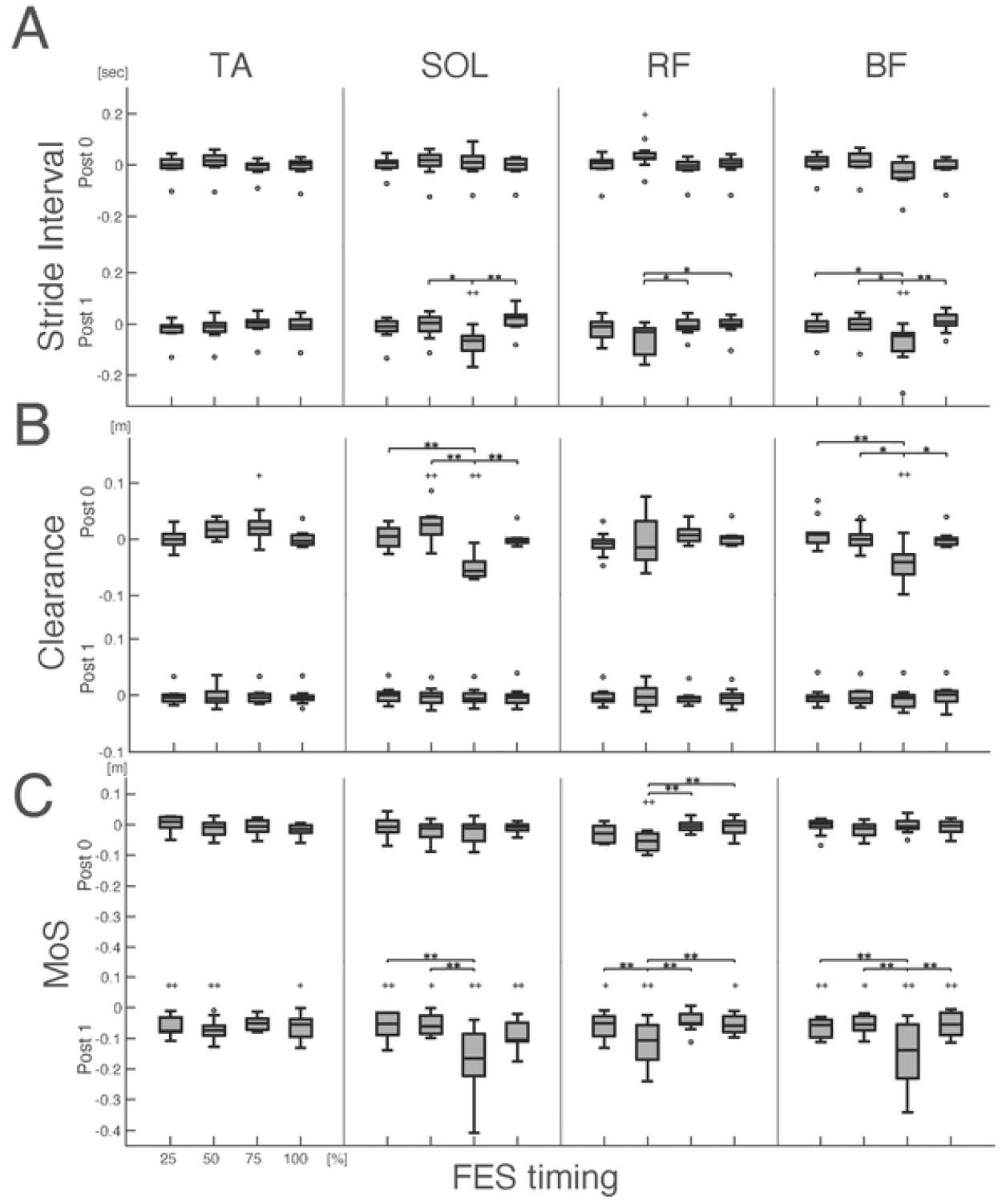
The displacement from the baseline in (A) Stride Interval, and (B) Clearance, and the largest negative displacement from the baseline in (C) MoS. The + symbol indicates a significant difference from the baseline (+: p < 0.05, ++: p < 0.01), while the * symbol indicates a significant difference between the ones at the timing conditions within each target muscle condition (*: p < 0.05, **: p < 0.01).

For foot clearance, significant positive displacements were found under TA75% and SOL50%, while significant negative displacements were observed under SOL75% and BF75% (Figure 3B). Among these, SOL75% and BF75% exhibited significantly lower clearances than other timing conditions. However, TA75% did not differ significantly from the other TA conditions.

For MoS, a significant reduction from baseline was observed at Post0 under RF50%, and across multiple timing conditions at Post1. All timing conditions under SOL and BF showed significant MoS decreases at Post1, with the greatest reductions occurring at SOL75%, RF50%, and BF75% (Figure 3C). In the SOL condition, the MoS reduction at 75% was larger than at the other timings, although not significantly different from 100%. No significant MoS differences were found among timing conditions within TA.

These three conditions—SOL75%, RF50%, and BF75%—were the most effective in reducing MoS. To directly compare these, Figure 4 presents stride interval (Figure 4A), clearance (Figure 4B), and MoS (Figure 4C) for these conditions.

**Figure 4.**
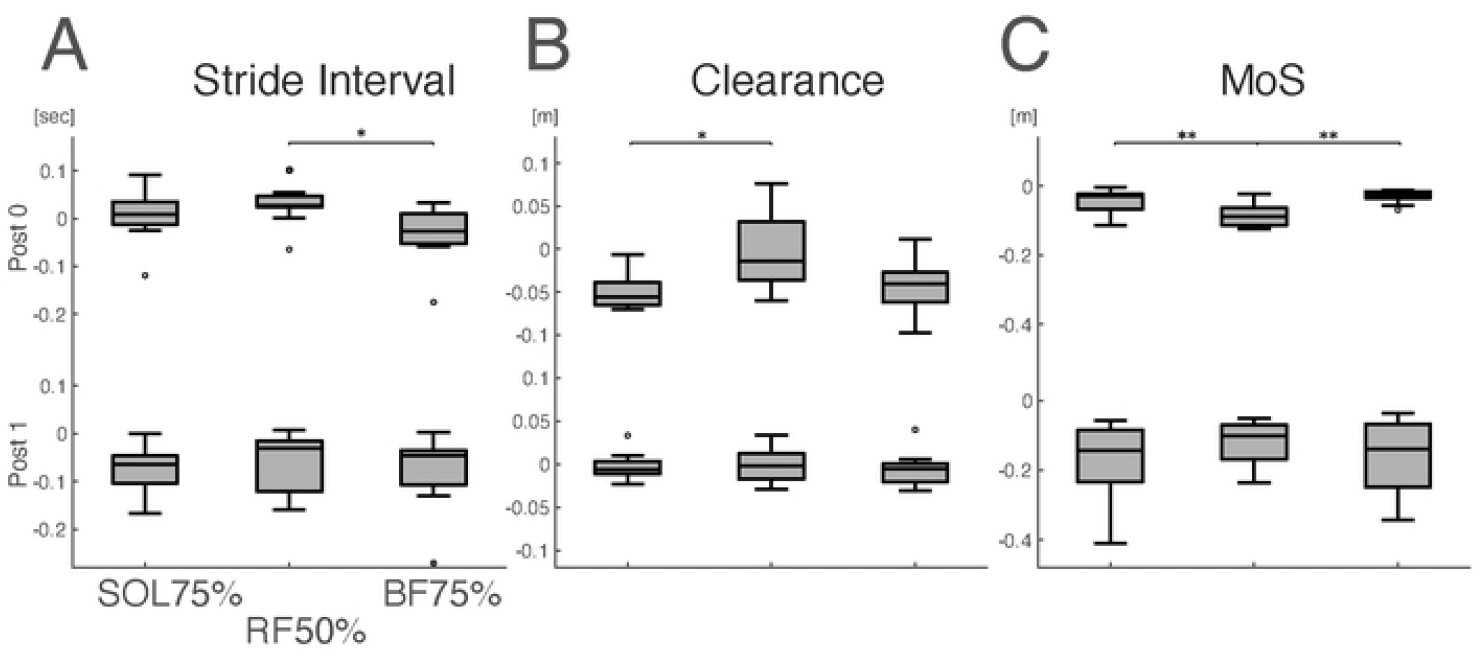
A comparison of (A) Stride Interval, (B) Clearance, and (C) MoS between the SOL75%, the RF50%, and the BF75%, which exhibited greater MoS decreases across all conditions. The * symbol indicates a significant difference between the ones at the timing conditions within each target muscle condition (*: p < 0.05, **: p < 0.01).

At Post0, only RF50% showed a significantly longer stride interval than baseline. However, when comparing across the three conditions, a significant difference was found only between RF50% and BF75%. This likely reflects higher inter-subject variability in stride interval under SOL75%.

For clearance, SOL75% and BF75% showed significant reductions from baseline, with a significant difference only between SOL75% and RF50%. This suggests that clearance was relatively unaffected under RF50%, while it tended to decrease under BF75%, with individual variation influencing the result.

For MoS, at Post0, only RF50% showed a significantly greater decrease from baseline compared to both SOL75% and BF75%. By Post1, all three conditions exhibited similarly reduced MoS values, with no significant differences between them. This indicates that while MoS reductions are evident in all three conditions by Post1, the effect emerges earlier under RF50%.

## Discussion

This study investigated the effects of DFES as an internal perturbation strategy to induce postural instability during treadmill walking. Among the 16 tested conditions, DFES applied to the soleus at 75% of the gait cycle (SOL75%), rectus femoris at 50% (RF50%), and biceps femoris at 75% (BF75%) resulted in the greatest decreases in the MoS. Notably, only RF50% produced a significant MoS reduction immediately after stimulation (Post0), indicating it was the most effective in provoking acute postural instability.

These DFES-induced instabilities were accompanied by atypical joint movements. In all three effective conditions (SOL75%, RF50%, BF75%), knee flexion at heel contact increased compared to baseline. Additionally, changes in ankle joint behavior, i.e., greater plantarflexion at toe-off or reduced dorsiflexion in late swing, were observed. These patterns are consistent with the known roles of the stimulated muscles: the SOL contributes to plantarflexion, the BF to knee flexion, and the RF to knee extension. For instance, the increased knee flexion under BF75% aligns with enhanced BF activity due to FES. Similarly, the reduction in dorsiflexion under SOL75% aligns with SOL activation.

Interestingly, greater knee flexion at heel contact under RF50% was somewhat counterintuitive, as the RF is typically associated with knee extension. However, the observed joint behavior at toe-off under RF50%, i.e., increased ankle plantarflexion and decreased knee flexion, suggests that the greater knee flexion at heel contact may have been a compensatory response to preceding atypical limb configurations. Similarly, the reduced ankle dorsiflexion observed under SOL75% in late swing may have contributed to the increased knee flexion at heel contact, reinforcing the interconnected role of the ankle and knee in impact absorption and postural control during walking [22–26].

In addition to joint kinematics, stride interval was significantly shortened under SOL75% and BF75%. Prior studies have shown that periodic stimulation of the calf and hamstring muscles can entrain the gait cycle and modulate stride timing [27,28]. Our findings support this, suggesting that entrainment or sensory modulation by DFES may alter temporal gait structure in ways that reduce dynamic stability. Together, these results indicate that both atypical joint mechanics and temporal disruptions may contribute to DFES-induced postural instability.

Notably, FES applied to the tibialis anterior (TA), i.e., a dorsiflexor active during early swing, did not lead to significant MoS reduction, despite inducing greater dorsiflexion at the ankle under TA50%. This suggests that DFES applied to muscles like the TA may not sufficiently disrupt postural control, possibly due to their lower contribution to load-bearing and impact phases of gait.

Our findings align with and extend the work of Manczurowsky et al. [15], who first introduced the concept of DFES to perturb internal motor coordination during walking. Their study focused on cognitive consequences, showing that DFES-induced discomfort may alter executive function. In contrast, our study systematically evaluated the motor consequences of DFES by manipulating stimulation timing and target muscle. While Manczurowsky et al. [15] selected stimulation parameters based on pilot testing, our findings provide a more comprehensive map of how DFES affects postural control, identifying specific conditions that reliably provoke instability. As noted by Budde and Gronwald [16], the DFES paradigm offers a valuable alternative to traditional mechanical perturbations by simulating internally generated impairments while avoiding confounds associated with clinical populations.

This study has several limitations. Although the walking speeds used span the reported preferred speeds for adults [17–21], individual differences in height (ranging from 160 to 180 cm) may have introduced variability, as preferred walking speed is influenced by leg length [29]. Additionally, FES timing was pre-programmed based on each participant’s stride interval and not dynamically adjusted in real time. Future studies should consider personalized or adaptive DFES protocols to enhance ecological validity.

In conclusion, this study demonstrated that DFES can successfully induce postural instability during walking. The conditions SOL75%, RF50%, and BF75% were particularly effective, with RF50% producing the most immediate disruption. These results highlight the potential of DFES as a tool to simulate internal motor impairments and investigate mechanisms of falls. By mimicking inadequate or mistimed muscle activity, DFES offers a promising avenue for developing more realistic fall models and testing targeted fall prevention strategies.

## Acknowledgements

We thank to Derrick Lim for his technical support during this study. This work was a supported by grants from the Natural Sciences and Engineering Research Council of Canada (grant no. DG RGPIN-2017-06790).

